# Multi-System Dysregulation in Placental Malaria Contributes to Adverse Perinatal Outcomes in Mice

**DOI:** 10.1101/2025.01.15.633265

**Authors:** Phebe Ekregbesi, Brittany Seibert, Maclaine A. Parish, Yevel Flores-Garcia, Patrick S. Creisher, Joseph P. Hoffmann, Jennifer Liu, Cory Brayton, Fidel Zavala, Sabra L. Klein

## Abstract

Sequestration of *Plasmodium* parasites in the placental vasculature causes increased morbidity and mortality in pregnant compared to non-pregnant patients in malaria-endemic regions. In this study, outbred pregnant CD1 mice with semi allogeneic fetuses were infected with transgenic *Plasmodium berghei* or mock-inoculated by mosquito bite at either embryonic day (E) 6 (first trimester-equivalent) or 10 (second trimester-equivalent) and compared with non-pregnant females. *P. berghei*-infected mosquitoes had greater biting avidity for E10 dams than uninfected mosquitoes, which was not apparent for E6 dams nor non-pregnant females. Infected E10 dams had greater numbers of parasites than E6 dams in the uterus and spleen, but not in the blood or liver. While parasites were found in placentas, no parasites were present in fetuses. Maternal infection at E6 caused greater maternal morbidity, with greater rates of fetal reabsorption and stillbirths than at E10. Infection at E10 caused adverse offspring outcomes, including growth restriction. To identify possible mechanisms of adverse offspring outcomes, E10 dams were euthanized during peak parasitemia (8 days post infection), and outcomes were compared with mock-infected dams. *P. berghei* caused significant systemic maternal immune activation with elevated circulating lymphocytes, eosinophils, and neutrophils and splenic cytokine concentrations. *P. berghei* infection at E10 increased corticosterone and decreased progesterone concentrations, which could contribute to adverse perinatal outcomes through immunomodulation. There were limited changes in the maternal fecal microbiome after *P. berghei* infection. Mosquito bite infection of outbred dams with *P. berghei* causes placental malaria and provides a novel, tractable model to investigate therapeutic treatments.

## INTRODUCTION

Malaria infection during pregnancy is a major public health threat worldwide. In 2020, 49.2% of pregnancies were at risk of malaria infection globally [1]. Pregnant people, especially first-time mothers, are at a higher risk of disease burden in incidence, frequency, and morbidity [2, 3]. Gestational malaria results in inflammation, coagulation, and placental tissue damage, causing fetal, neonatal, and maternal mortality [4]. While the mechanisms of gestational malaria are unknown, disruption to the placenta by *Plasmodium* parasites is associated with intrauterine growth restriction, preterm labor, miscarriage, and preeclampsia [3]. Further, maternal malaria-induced fetal growth restriction is estimated to have a mortality rate of 37.5% each year [5]. Malaria infection in pregnant patients can develop into placental malaria [6], which is the sequestration of *Plasmodium* parasites in the intervillous space of the placenta, often without peripheral parasitemia [4]. Clinically, placental malaria is difficult to diagnose as parasite sequestration in the placenta cannot be detected by microscopy or rapid diagnostic tests that use peripheral samples [4, 7]. Histopathology is the gold standard for placental malaria diagnosis, but it can only be performed post-partum [4]. Directly monitoring placental malaria is challenging; therefore, systemic *Plasmodium* infection is monitored, which, although easier to assess and a prerequisite of placental malaria, can often underestimate the condition. The more apparent fetal outcomes are prioritized, with interventions often focused on protecting the child, with maternal outcomes often neglected. Investigating the mechanisms of placental malaria pathogenesis is fundamental for improving maternal and child outcomes.

While numerous factors can impact susceptibility to placental malaria and malaria infection during pregnancy, placental insufficiency is often the underlying mechanism of many obstetric complications [8, 9]. During placental malaria, infected RBCs (iRBCs) accumulate in the intervillous space of the placenta, inducing syncytial knots and fibrin deposition [4]. As iRBCs lyse, hemozoin and oxidizing free heme are released, signaling macrophage infiltration into the intervillous space [4]. VAR2CSA- mediated cytoadherence initiates an inflammatory cascade, which is exacerbated as parasites replicate and evade clearance [10]. Proinflammatory cytokines, oxidative stress, and increased cell death can result in placental pathology and disrupted transplacental nutrient exchange, ultimately resulting in poor birth outcomes [4]. Further research into the mechanisms of placental malaria pathogenesis is essential for developing additional treatments to mitigate severe disease in pregnant patients and improve birth outcomes.

*Plasmodium* parasites, the causative agent of malaria, have a broad host range and are transmitted by female *Anopheline* mosquitoes. In humans, the five *Plasmodium* species that cause malaria are *Plasmodium falciparum, Plasmodium vivax, Plasmodium ovale, Plasmodium malariae*, and *Plasmodium knowlesi* [11]. Etiologic species in rodents include *Plasmodium berghei, Plasmodium chabaudi, Plasmodium yoelii,* and *Plasmodium vinckei* [12]. *P. berghei* and *P. chabaudi* are the most extensively used parasites in placental malaria mouse models. *P. berghei* blood-stage pathogenesis closely resembles *P. vivax* [12]. Similar to *P*. *vivax*, *P. berghei* merozoites target reticulocytes and lack synchronicity [12]. In naïve mouse models of placental malaria, infection is typically achieved via intraperitoneal (i.p.) injection of a high dose of iRBCs that induce rapid mortality [13, 14]. In rare cases, infection is performed through intravenous (i.v.) injection of sporozoites into inbred mice [13, 14]. While inbred mouse strains are preferred in placental malaria models for their genetic tractability, syngeneic pregnancies in inbred mice do not model the maternal-fetal immune tolerance needed to preserve a genetically distinct fetus in allogenic pregnancies [15, 16]. In addition to inbred mice, knock-out mice have been used to investigate targeted mechanisms of placental malaria pathogenesis. Recent mouse placental malaria models have begun utilizing outbred mice [17, 18] or crossbreeding different inbred mouse strains [19] to simulate allogenic pregnancies.

Mosquito bite challenge, a more physiologically relevant method of inoculation, may delay lethality compared to current models, providing a useful tool for studying infection during pregnancy and allowing sufficient survival to evaluate birth outcomes. Thus, we developed a translational placental malaria model using outbred CD-1 mice to simulate semi-allogeneic maternal-fetal tolerance and employ mosquito bite challenge for low-dose sporozoite inoculation. We utilized this model to investigate whether placental malaria induces maternal immune, endocrine, and microbiome changes to drive poor birth outcomes. By developing a new physiologically relevant model of placental malaria, the timing of malaria infection on pathogenesis can be interrogated. This model could serve as a preclinical model to test therapeutics, such as monoclonal antibodies and vaccines, to mitigate the impact of malaria on pregnancy outcomes.

## MATERIALS & METHODS

### Parasites and Mosquitoes

*Plasmodium berghei*, Strain (ANKA) 676m1cl1, MRA-868, was obtained through BEI Resources, NIAID, NIH, and contributed by Chris J. Janse and Andrew P. Waters. The transgenic *P. berghei* ANKA reporter line expresses GFP-luciferase throughout the parasite life cycle and does not contain a drug-selectable marker. Parasites were amplified in female Swiss-Webster mice by the Johns Hopkins Malaria Research Institute Parasite core. *P. berghei* GFP-luc (MRA-868) parasites were used in all experiments. *Anopheles stephensi* mosquitoes were provided by the Johns Hopkins Malaria Research Institute Insectary. Four-day old mosquitoes were infected by feeding on Swiss Webster mice infected with *P. berghei* GFP-luc. After infection, mosquitoes were maintained in an incubator. Mosquito inoculation was confirmed by microscopy at the oocyst and sporozoite stage at 14 and 21 days post-feed, respectively [20]. For mock inoculations, a separate cage of uninfected mosquitoes was used. All mosquitoes were supplied with 10% sucrose except for 24 hours prior to bloodmeals.

### Mouse experiments

Timed-pregnant CD-1 IGS dams were ordered at embryonic day (E)4 or E8 from Charles River Laboratories (Wilmington, MA) and single-housed throughout the study until delivery or death. Mice were randomly assigned by weight to either the *Pb*- or mock-inoculated groups 1 day after arrival. At E6 or E10, dams were anesthetized with 0.5 ml of 2% Avertin intraperitoneally and exposed to 5 female *Pb*-infected *Anopheles stephensi* mosquitoes per mouse. The number of infected female mosquitoes used for infections was calculated based on the cage sporozoite prevalence rate. Mock-inoculated dams were exposed to uninfected female mosquitoes. Blood-feeding avidity was calculated for each dam as the percentage of engorged female mosquitoes divided by the total number of female mosquitoes. After infection, body mass, rectal temperature, and clinical signs were recorded daily to monitor morbidity until mice were euthanized or succumbed to infection [21]. Thin blood smears from tail incisions were collected prior to infection and daily from 3-21 days post-infection (dpi), and stained with Giemsa stain (Sigma, St. Louis, MO) to monitor peripheral parasitemia. All animal experimental procedures and outcomes were approved by the Johns Hopkins Animal Care and Use Committee (MO21H246).

### Tissue and Serum Collection

Plasma was collected from tail incisions at 8 dpi. A subset of dams was euthanized at 8 dpi using 160 mg/kg ketamine and 10 mg/kg xylazine solution. Upon euthanasia, livers, spleens, uteri, placentas, and fetuses were collected and used for intra-vital imaging system (IVIS) imaging. Snap-frozen placentas and spleens were homogenized, and cytokine multiplexing was performed on tissue homogenates. For histological analysis, one uterine horn was fixed in 10% neutral buffered formalin (Sigma, St. Louis, MO).

### Pregnancy Outcomes

Pregnancy was confirmed by laparotomy upon maternal death, euthanasia, or by spontaneous delivery. Pregnancy outcomes were determined within 12 hours of birth or dam death. In dams followed until death, newborn mass, resorptions, viability, and stillbirths were recorded as outlined [17, 19]. The mass of each viable pup was taken at postnatal day (PND) 0 and recorded as an average of the total viable litter size. Black compact footprints on the uterine horn were considered resorptions. Pink, well-vascularized fetuses were considered viable, and resorbed or grey/discolored embryos were considered non-viable. Dams with all non-viable fetuses were recorded as a pregnancy loss and expressed as a percentage of the total number of dams surviving to or beyond E19 per group. The birth of dead fetuses or fetuses attached to the placenta were considered stillbirths. Stillbirths were recorded as a percentage of the total litter size (including non-viable pups). In dams euthanized at 8 dpi, placental efficiency and fetal viability were assessed. The fetus, placenta, and uterine wall from implantation sites were dissected for further analysis. Fetal viability was assessed as previously described [22], and non-viable pups were counted and recorded as a percentage of total implantation sites.

### Parasite Quantification

Peripheral parasitemia was assessed by counting iRBCs in blood smears. Three non-consecutive fields were imaged on the Eclipse E200 (Nikon, Tokyo, Japan) microscope. All RBCs within a 10×10 reticle were counted using the Cell Counter plugin in ImageJ software (version 1.54c) [23]. Parasitemia was defined as infected RBCs calculated as a percentage of total RBC.

### Intra-Vital Imaging System Analyses (IVIS)

Tissue parasite burden was determined by intra-vital imaging system (IVIS). Briefly, dams were anesthetized with 160 mg/kg ketamine and 10 mg/kg xylazine solution IP. When mice were non-responsive to toe-pinch, they were administered 100 μL of 30 mg/mL D-luciferin (Perkin Elmer, Waltham, MA) retro-orbitally. After 2 minutes, deeply anesthetized mice were bled and humanely euthanized. Dams were immediately dissected to remove the liver, spleen, uterus, placentas, and fetuses from 3 implantation sites. Tissues were imaged on the IVIS Spectrum (Perkin Elmer, Waltham, MA) and the Living Image software (version 4.7.4, Perkin Elmer, Waltham, MA) in the Luminescent mode. Exposure time was set at 20 seconds, with a 1-minute delay, medium binning, and F/stop 1. Bioluminescence was recorded as average radiance measured as photons/second/centimeter^2^/steradian.

### Histology

Individual placentas were dissected from uterine horns and fixed in 10% neutral buffered formalin (Sigma, St. Louis, MO) until tissues were processed for staining by the Johns Hopkins Reference Histology Laboratory. H&E staining was used to determine morphological changes in placentas. Giemsa staining was used to identify infected red blood cell cytoadherence to the placental endothelium. H&E-stained slides were imaged using an Eclipse 80i (Nikon, Tokyo, Japan) at 2x, 10x, and 40x magnifications. Micrographs of five non-consecutive fields were recorded digitally using CaptaVision software version 2.4.1 (AccuScope, Commack, NY). Placentas were scored for inflammation, cell death, and fibrin deposition by an ACVP certified veterinary pathologist (CB). Giemsa-stained slides were imaged using an Eclipse E200 (Nikon, Tokyo, Japan) at 10x and 100X magnifications. Qualitative image analysis was conducted using ImageJ (version 1.54c) [23].

### Hematology

Blood (100μL) from terminal cardiocenteses was collected into EDTA tubes. Automated complete blood counts (CBC) were performed within 6h on an ProCyte Dx (Idexx) hematology analyzer.

### Multiplex Cytokine Analysis

Cytokine and chemokine concentrations in spleen and placental homogenate were measured using the Procartaplex Mouse Immune Monitoring 48-plex Panel (Invitrogen, Waltham, MA) following standard protocol. Briefly, serially diluted standard or neat tissue homogenate samples were incubated with capture beads before antibody detection, followed by indirect staining with streptavidin-PE. Beads were resuspended in assay buffer and then read on an xMAP Intelliflex System (Luminex, Austin, TX). Analysis was performed using the ProcartaPlex™ Analysis App on Thermo Fisher Connect and visualized using GraphPad Prism version 10.1.2 (GraphPad Software, Boston, MA).

### Steroid Hormone ELISAs

Progesterone ELISAs were performed as previously described [24]. Briefly, steroids were extracted from 10 μL tail bleed sera in diethyl ether in a 1:10 (tail sera) or 1:5 (placental homogenate) sample-to-ether ratio. Extracted steroids were further diluted 1:5 (tail sera) or 1:10 (placental homogenate) in assay buffer and progesterone was quantified by ELISA following the manufacturer’s instructions (Enzo Life Sciences, Long Island, NY). Plates were read at 405 nm in a SoftMaxPro5 plate reader, and concentrations were determined from a 4PL standard curve generated in GraphPad Prism version 10.1.2 (GraphPad Software, Boston, MA). To measure corticosterone, steroid extract from the tail was diluted 1:100 in assay buffer before corticosterone was quantified by ELISA according to the manufacturer’s instructions (Arbor Assay, Ann Arbor, MI). Plates were read at 450 nm in a SoftMaxPro5 plate reader, and concentrations were determined from a 4PL standard curve generated in SoftMaxPro software version 7.1 (GraphPad Software, Boston, MA). All samples analyzed for progesterone and corticosterone were assayed in duplicate.

### DNA extraction and shotgun metagenomic sequencing library preparation

DNA was extracted from free-drop fecal samples collected at 8 dpi using the Qiagen DNeasy PowerLyzer PowerSoil kit (Qiagen, Gaithersburg, MD). Following extraction, 1 ng of DNA was used in the Nextera XT Library preparation workflow. Genomic DNA was initially tagmented, after which dual indexes and terminal complimentary flow cell oligos were added through a PCR reaction following the Nextera PCR protocol. Libraries were cleaned using AMPure XP beads (Beckman Coulter, Indianapolis, IN) at a 3:2 ratio. Libraries were checked on the Agilent High Sensitivity D5000 ScreenTape (Santa Clara, CA). Normalized libraries were pooled, diluted, and sequenced using the Illumina NovaSeq X Plus 10B (PE150) platform (Novogene, China). Negative controls, including an extraction blank, were included.

### Sequence processing and analysis

Primer removal and demultiplexing were performed by Novogene using Illumina BaseSpace software. The quality of the raw paired-end reads was visualized using FastQC. Read 2 was trimmed by 20 base pairs (bp) and reads with an average quality below 20 or a total length shorter than 100 bp were discarded using BBduk from the BBtools software [25]. Host sequences were removed using BBsplit [25] with default settings and the GRCm39 mouse genome reference. Taxonomy was assigned using Metaphlan4 [26] with default settings, which was used for downstream analysis.

Alpha (Richness and Shannon diversity) and beta diversity (Bray-Curtis dissimilarity) metrics were calculated using Metaphlan4. Results were imported into the *phyloseq* [27] package using R Statistical Software (v4.4.1; [28]). An ordination with a pre-computed distance matrix was performed using “ordinate” from *phyloseq*. Results were graphed using *ggplot2* [29] and GraphPad Prism version 10.1.2 (GraphPad Software, Boston, MA). All statistical tests were performed using multiple Mann-Whitney tests. Multivariate statistics analysis was conducted using permutational multivariate analysis of variance (PERMANOVA) based on Bray-Curtis dissimilarity distances with 1,000 permutations generated via the *vegan* [30] package’s “adonis2” function. For taxonomic analysis, samples were first filtered to remove reads that did not match the kingdom Bacteria. Relative abundances at the phylum and species levels were calculated by aggregating data based on infection status, pregnancy status, and taxonomic level. Taxonomic bar plots were generated using *ggplot2,* representing species greater than 2% relative abundance. Differential analysis comparing mock and *Pb*-infected E10 dams was performed using linear discriminant analysis effect size (LEfSe) [31]. A p-value <0.05 was considered significant. Correlation analysis was performed to better understand the relationship between taxa and different host factors. Spearman correlation analysis was calculated with relative abundance data, and a simple linear regression analysis of the correlation data was performed.

### Statistical Analyses

Statistical analyses were performed using GraphPad Prism version 10.1.2 (GraphPad Software, Boston, MA). Comparisons of three or more groups by gestational age, infection status, and/or organ for blood-feeding avidity, parasite radiance, and serum progesterone were done using two-way ANOVA with Tukey correction. Kaplan-Meier curves for each gestational age were compared using the log-rank Mantel-Cox test. Differences between parasitemia curves across the gestational ages were assessed by a one-way ANOVA with Tukey post-hoc test. For cytokine concentration, complete blood count, and steroid concentrations, comparisons between mock- and *P. berghei*-infected E10 dams were performed using unpaired Student’s t-test. A simple linear regression analysis of the correlation data was performed. For dams that were followed until they delivered or succumbed to infection, differences in offspring outcomes (embryonic day of delivery, pregnancy loss, and stillbirths) by infection status were compared using Fisher’s exact test for each gestational age. Differences in fetal outcomes (viability, fetal and placental mass, and placental efficiency) for offspring of E10 dams, collected at 8 dpi, by infection status were compared using an unpaired Student’s t-test. Significance was set at p < 0.05 and is indicated by an asterisk for graphs or bolded text for tables.

## RESULTS

### Maternal *Pb* challenge during either the first or second trimester causes placental infection

Non-pregnant and pregnant CD-1 mice were exposed to mock (uninfected)- or *Pb*-infected female mosquitoes, with tissues collected 8 days post-infection (dpi) (**Fig 1A**). While previous placental malaria models infect at embryonic day (E) 12-14 due to the high lethality of blood stage infection, we exposed pregnant dams at earlier gestational ages, i.e., E6 or E10, to examine *Pb* exposure during the first and second trimester-equivalent, respectively. To determine if pregnancy status, gestational age, or both could impact the mosquitoes’ feeding behavior, blood-feeding avidity was analyzed. While mock and *Pb*-infected mosquitos showed similar preference for non-pregnant and E6 dams, blood-feeding avidity was significantly greater among *Pb*- infected mosquitoes compared to mock-infected mosquitoes exposed to E10 dams **(Figure 1B).**

**Figure 1.**
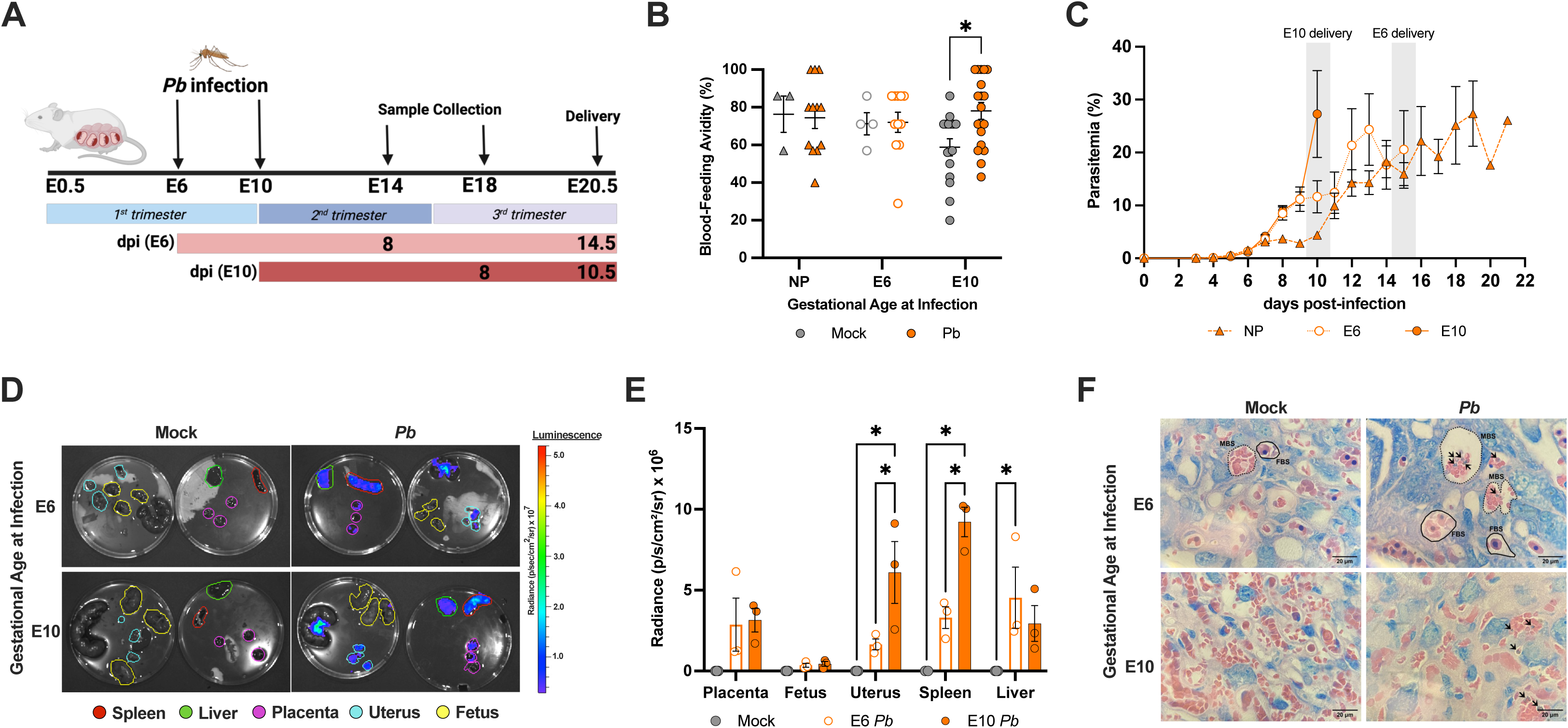
Mosquito bite *Plasmodium berghei* challenge results in infection within the placenta. **(A)** CD-1 dams were infected on embryonic day (E)6 or E10 by mosquitos infected with *P. berghei* (*Pb*) GFP-luc and monitored until they succumbed to infection or delivered. A subset of dams was infected on E6 or E10 and followed for 8 days post-infection (dpi), after which they were euthanized, and tissues were collected. **(B)** Blood-feeding avidity of mock- and *Pb*-infected female mosquitoes during blood meals for non-pregnant (NP) females (triangles) and pregnant dams infected at E6 (open circles) or E10 (closed circles). n = 3-18 dams/group across 6 experiments. **(C)** Parasitemia was measured by blood smear with Giemsa stain of non-pregnant or pregnant dams infected at E6 or E10. n = 12-18 dams/group across 6 experiments. **(D)** Representative luminescent images of the spleen, liver, placenta, uterus, and fetus (differentiated by the outline color) from dams infected at E6 or E10, captured by the IVIS at 8 dpi. **(E)** Tissue parasite burden quantified by luciferase radiance from the IVIS in tissues collected at 8 dpi. n = 3 dams/group. **(F)** Representative histology of Giemsa-stained placental sections from E6 and E10 dams collected at 8 dpi, labyrinth zone imaged at 100X magnification (scale bar, 20 μm). Infected red blood cells (iRBC) are indicated by arrows. Dotted outlines indicate maternal blood spaces (MBS), and solid outlines indicate fetal blood spaces (FBS). Data was analyzed using a 2-way ANOVA with Tukey post-hoc test. Asterisks indicate p < 0.05.

Parasitemia was evaluated from blood smears collected on the day of infection and every day from 3 days post-infection (dpi) until death. Parasitemia was first detected in mice at 5 dpi in pregnant and non-pregnant mice; however, pregnant mice, especially those in the E10-infected group, exhibited a faster onset of peak parasite burden than non-pregnant mice (**Fig 1C**). In our model, mosquito bite infection delayed the onset of parasitemia compared to malaria models utilizing injection of iRBC [32–35]. To evaluate tissue penetration of malaria parasites following mosquito bite challenge, the parasite burden in tissue vasculature was analyzed by bioluminescence after infection with *P. berghei* GFP-luc using IVIS imaging. Among dams infected at either E6 or E10, the luminescent signal was detected in all maternal tissues, including the liver, spleen, and uterus, and was significantly higher in the uteri and spleens from *Pb*- infected E10 than in E6 dams (**Figure 1D**). Luminescent signal was detected in the placentas and liver of both E6 and E10-infected dams, but no luminescent signal was detected in the fetuses from dams infected with *Pb* at either E6 or E10 (**Figure 1D and 1E**). To further confirm *Pb* infection of placental tissue, Giemsa staining was used to visualize parasites in tissue. Placentas from dams infected at either E6 or E10 had iRBCs in the maternal intervillous spaces of the placenta (**Figure 1F**). Taken together, these data suggest that mosquito-bite *Pb* infection during pregnancy results in efficient maternal parasite dissemination, including to the placenta at 8 dpi.

### Gestational age impacts morbidity and mortality in dams and offspring

With *Pb* infecting multiple tissues, including the placenta, we next examined maternal morbidity as well as fetal and perinatal outcomes following *Pb* infection at E6 or E10. E6-infected dams gained less body mass during the third trimester of pregnancy (E16-21) than their mock-infected controls (**Figure 2A**). In contrast, E10-infected dams had body mass gain comparable to mock-infected dams. *Pb* infection by mosquito bite challenge was lethal in our model within 9 -15 dpi for pregnant dams and 8-22 dpi in non-pregnant females **(Supplemental Figure 1A)**. *Pb* infection was uniformly lethal for E6 and E10-infected dams; however, mortality was first observed at E15 in E6-infected dams, whereas it occurred at E19 in E10-infected dams (**Figure 2B**). This suggests that in our model, morbidity and mortality prior to delivery are more pronounced in dams exposed to *Pb* earlier in pregnancy.

**Figure 2.**
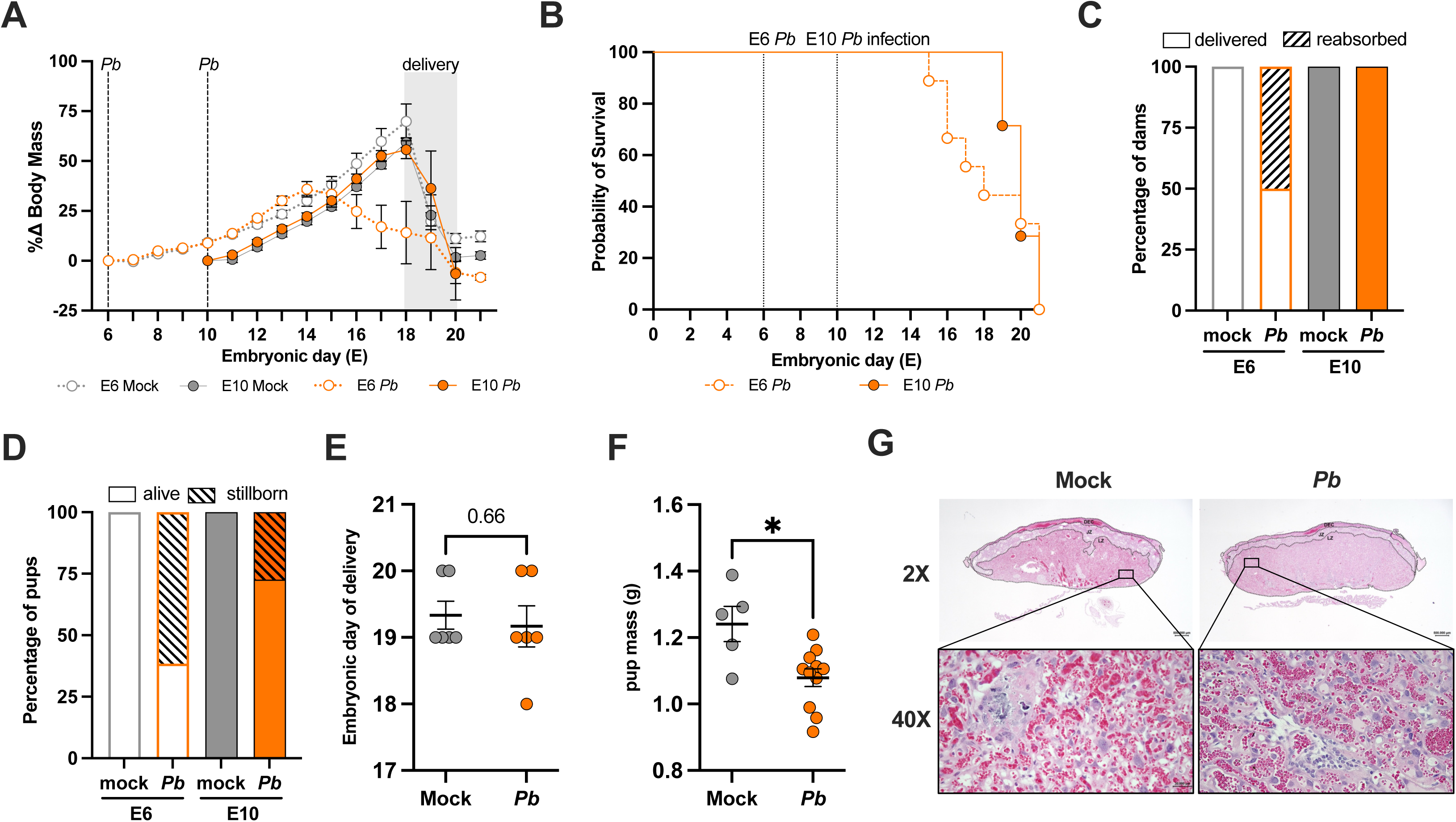
*P. berghei* infection during pregnancy causes maternal and perinatal morbidity and mortality in a gestational age-dependent manner. (A) Body mass change was calculated for dams infected at either embryonic day (E) 6 or E10 across embryonic days. Black stippled lines indicate the day of *P. berghei* (*Pb*) infection at E6 or E10 or the average day of delivery. n = 3-18 dams/group across 6 experiments. **(B)** Kaplan-Meier survival curves for E6 and E10 *Pb*-infected dams across embryonic days. n = 7-12 dams/group across 6 experiments. **(C)** The percentage of dams with reabsorption was calculated as the percentage of pups that survived until E19. Data was analyzed across 2 experiments. **(D)** The percentage of stillborn pups was calculated as a percentage of the number of stillborn pups divided by the total litter size. Data was analyzed across 2 experiments**. (E)** Embryonic day of delivery of mock or *Pb*- infected E10 dams. **(F)** Mass of fetuses from mock or *Pb*-infected E10 dams collected by cesarian section at 8 days post-infection. Statistical analysis was performed using an unpaired t-test. n= 5-11 pups/group across 2 experiments. Asterisks indicate p < 0.05. **(G)** Representative histology of H&E-stained placental sections from E10 dams collected at 8 dpi. Labyrinth zone imaged at 2X magnification (scale bar, 500 μm) and 40X magnification (scale bar, 30 μm).

Because adverse pregnancy outcomes can be a consequence of placental malaria, we analyzed adverse fetal outcomes by assessing the percentage of reabsorption and stillborn pups in dams that survived to at least E19 (i.e., the average embryonic day of delivery). The percentage of reabsorption was measured as the proportion of dams reabsorbing at least 1 conceptus, while the percentage of stillbirths was measured as the number of pups stillborn relative to the number of total pups within a litter. Reabsorption was observed in E6- but not E10-infected dams, with 50% of *Pb*- infected E6 dams experiencing pregnancy loss (**Figure 2C**). Stillbirths were associated with *Pb* infection at both E6 and E10, with 61.5% and 27.4% of pups from *Pb*-infected E6 and E10 dams, respectively, being stillborn (**Figure 2D**). These data suggest that gestational age affects malaria outcome, with infection during pregnancy being more detrimental during the first (E6) than second (E10) trimester-equivalent of pregnancy in our model.

Because E10-infected dams experienced slower mortality combined with no evidence of offspring reabsorption or stillbirths compared to E6-infected dams, subsequent analyses of neonatal outcomes were performed in E10-infected dams only. Fetal viability at E18 was analyzed by performing a cesarian section in E10 dams, with no significant difference between mock- and *Pb*-infected dams **(Supplemental Figure 1B)**. This observation suggests that stillbirths only occurred at or around the time of labor and delivery. The average embryonic day of delivery also did not differ between mock- and *Pb*-infected E10 dams, suggesting that *Pb*-infection at E10 did not induce preterm birth (**Figure 2E**). After birth, pups born from *Pb*-infected E10 dams had significantly decreased mass, suggesting that malaria during pregnancy causes intrauterine growth restriction in our model (**Figure 2F**). There was, however, no histological evidence of placental damage observed at 8 dpi (peak parasitemia) among dams that were either mock- or *Pb*-infected at E10 (**Figure 2G**).

### *Pb* infection during pregnancy causes maternal immune activation

Pregnancy requires an immunologically tolerant environment to support intrauterine growth of the fetus while also maintaining the ability to respond to potential maternal infections. We examined the immunophenotype of *Pb*-infected E10 dams at peak parasitemia (8 dpi). Complete blood counts were used to determine the overall clinical phenotype of infected dams (**Table 1**). Anemia was not observed by hematocrit or hemoglobin measures among dams infected with *Pb* at E10 (**Table 1**). Reticulocytes and circulating white blood cells were significantly elevated in *Pb*-infected compared with mock-infected E10 dams (**Table 1**). The increase in circulating white blood cells was driven by the significantly increased number of circulating lymphocytes (**Figure 3A**), eosinophils (**Figure 3B**), and neutrophils (**Figure 3C**). While neutrophils increased as a proportion of total leukocytes in response to infection, the lymphocyte fraction dropped, resulting in a significantly increased neutrophil-to-lymphocyte ratio (a measure of chronic stress [36]) in *Pb*-infected dams **(Figure 3D)**. Overall, malaria infection during pregnancy results in increased circulating white blood cells, suggesting increased systemic inflammation at peak parasitemia within our model.

**Figure 3.**
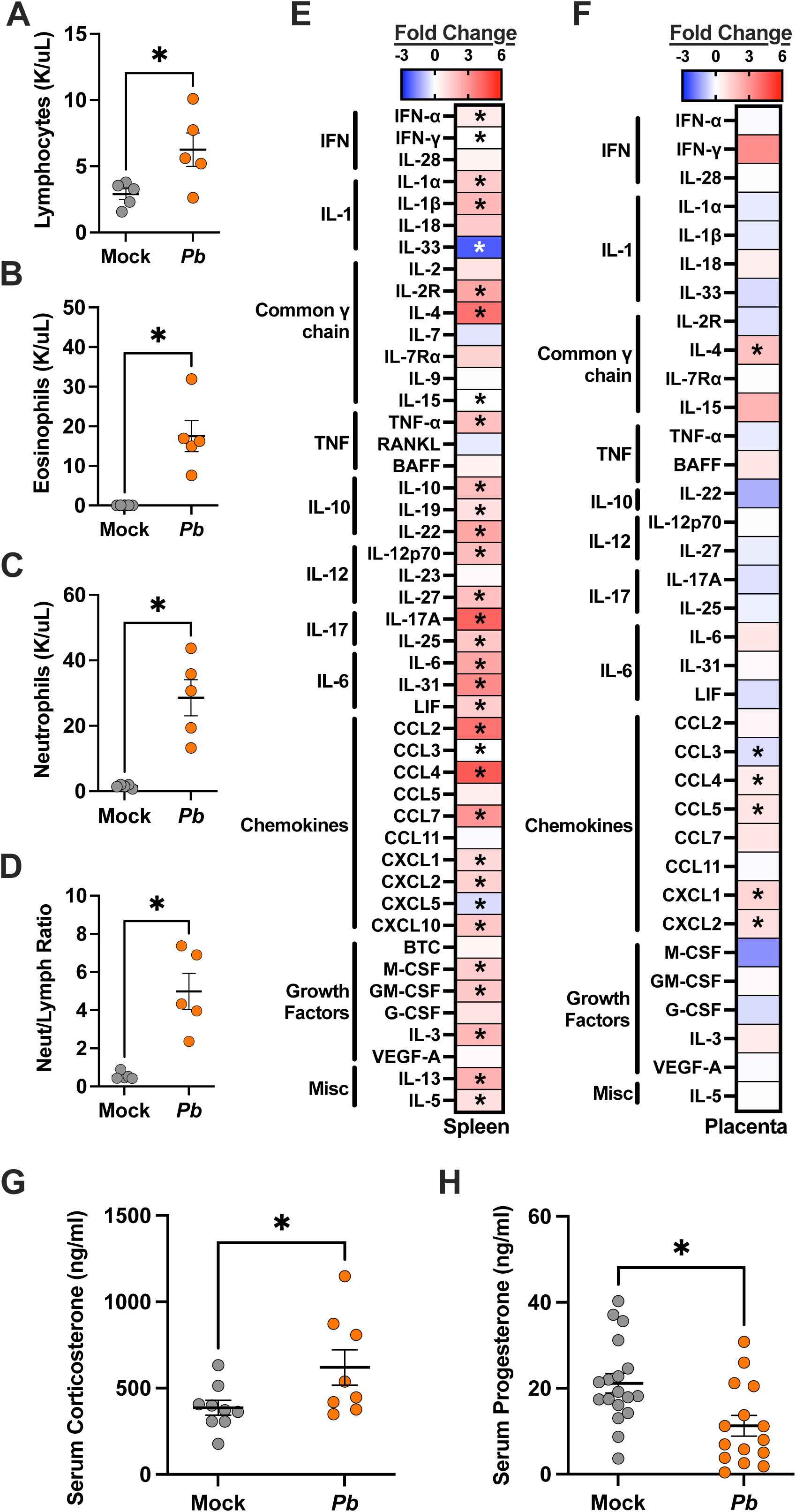
Malaria infection during pregnancy induces systemic inflammation and placental chemokine activation. **(A)** Lymphocyte, **(B)** eosinophil, **(C)** neutrophil, and **(D)** neutrophil/lymphocyte concentrations were analyzed from complete blood counts from E10 mock and *Pb*-infected dams at 8 dpi. Statistical analysis was performed using an unpaired t-test. n=5-6 per group across 2 experiments. A heatmap illustrating the log fold change in splenic **(E)** and placental **(F)** cytokine concentrations of *Pb*-infected E10 dams relative to the mock controls at 8 dpi n=5-9 per group across 2 experiments. **(G)** Corticosterone and **(H)** progesterone concentrations were measured in plasma from E10 dams at 8 days post-infection by ELISA. n = 8-16 dams/group across 4 experiments. Asterisks indicate p < 0.05.

**Table 1.** Complete blood counts for mock- and *Plasmodium berghei* (*Pb*)-infected E10 dams.

| Readout | Mock |  | *Pb* |  | *p*-value |
| --- | --- | --- | --- | --- | --- |
|  | Mean | SEM | Mean | SEM |  |
| Red blood cells (M/uL) | 7.24 | 1.52 | 8.17 | 0.62 | 0.612 |
| Hematocrit (%) | 38.20 | 7.97 | 42.98 | 3.59 | 0.623 |
| Mean cell volume (fL) | 52.40 | 1.48 | 52.50 | 1.24 | 0.961 |
| RDW-CV (%) | 23.74 | 0.77 | 23.00 | 0.76 | 0.511 |
| Hemoglobin (g/dL) | 11.88 | 2.44 | 13.94 | 1.09 | 0.493 |
| MCH (pg) | 13.75 | 2.76 | 17.04 | 0.23 | 0.311 |
| MCHC (g/dL) | 26.10 | 5.30 | 32.50 | 0.42 | 0.304 |
| **Reticulocytes (K/uL)** | **218.10** | **58.98** | **613.42** | **141.33** | **0.022** |
| **Reticulocytes (%)** | **3.06** | **0.67** | **7.25** | **1.23** | **0.017** |
| Immature reticulocyte fraction (%) | 41.06 | 10.30 | 40.98 | 7.08 | 0.995 |
| Platelets (K/uL) | 1232.17 | 755.41 | 297.00 | 18.92 | 0.292 |
| **White blood cells (K/uL)** | **4.06** | **0.89** | **53.08** | **8.62** | **0.000** |
| **Neutrophils (K/uL)** | **1.54** | **0.24** | **28.58** | **5.50** | **0.001** |
| **Neutrophils (% WBC)** | **31.32** | **2.66** | **53.36** | **4.92** | **0.004** |
| **Lymphocytes (K/uL)** | **2.90** | **0.42** | **6.26** | **1.26** | **0.035** |
| **Lymphocytes (% WBC)** | **59.54** | **3.77** | **11.94** | **1.71** | **0.000** |
| Monocytes (K/uL) | 0.35 | 0.08 | 0.61 | 0.18 | 0.207 |
| **Monocytes (% WBC)** | **8.14** | **2.43** | **1.58** | **0.49** | **0.029** |
| **Eosinophils (K/uL)** | **0.04** | **0.01** | **17.55** | **3.96** | **0.002** |
| **Eosinophils (% WBC)** | **0.88** | **0.21** | **33.10** | **4.84** | **0.000** |
| Basophils (K/uL) | 0.01 | 0.00 | 0.01 | 0.01 | >0.999 |
| Basophils (% WBC) | 0.12 | 0.05 | 0.02 | 0.02 | 0.095 |

E10 dams euthanized at 8 dpi, n=5-6 per group, data from 2 replicates. Bold indicates a statistically significant difference between mock and *Plasmodium berghei* groups, based on unpaired t-test, p-value <0.05. RDW-CV, red cell distribution width coefficient of variation; MCH, mean corpuscular hemoglobin; MCHC, mean corpuscular hemoglobin concentration; WBC, white blood cells; NE:LY ratio, neutrophil-to-lymphocyte ratio.

To further interrogate immune changes associated with maternal malaria, concentrations of 48 cytokines and chemokines were measured in spleens and placentas collected at 8 dpi from dams that were either mock or *Pb* infected at E10. Of the 46 cytokines and chemokines detected in the spleen, 33 (71%) were differentially expressed in the spleens of *Pb* and mock-infected dams (**Figure 3E**). Of the 33 analytes that were differentially expressed, 31 (94%) were significantly upregulated, and 2 (6%) were significantly downregulated in the spleens of *Pb-* compared with mock-infected dams (**Supplementary Table 1**). Splenic concentrations of proinflammatory mediators, such as type I and type II interferons, TNF-α, and cytokines from the IL-1 family and IL-6 family were significantly increased in *Pb-*infected dams. Traditional Th2 cytokines and members of the IL-17 family also were elevated in response to *Pb* infection. Chemokines, as well as hematopoietic, lymphocyte, and myeloid cell growth factors, increased in response to *Pb* infection. By comparison, the alarmin IL-33, which has been shown to be associated with protection from complicated malaria [37], was significantly decreased in the spleens of *Pb*-infected dams. Because placental infection, stillbirths, and intrauterine growth restriction were observed in E10-infected dams, we next investigated cytokines and chemokine concentrations in the placenta. Of the 48 analytes tested, 35 were detectable in placentas collected 8 dpi from mock- and *Pb*- infected dams. Of the 35 detectable analytes in the placenta, only 6 (17%) were differentially expressed between mock- and *Pb*-infected dams, 5 (83%) of which were significantly upregulated, and one (17%) was downregulated in placentas of *Pb*- compared with mock-infected dams (**Figure 3F** and **Supplementary Table 1**). In addition to IL-4, neutrophil and CD4+ T cell chemoattractants (CXCL1, CXCL2, CCL4, and CCL5) were significantly increased in the placentas of *Pb*-infected dams compared to mock-infected dams. In contrast, CCL3, which is associated with eosinophil recruitment and neutrophil activation, was significantly reduced in the placentas of *Pb*- infected dams compared to mock controls. Consistent with histological analyses, these results indicate that mosquito bite infection with *Pb* during mid-gestation does not cause large-scale placental inflammation at 8 dpi. The elevated chemoattractants of neutrophils and T cells in the presence of malaria parasites in the placentas and pregnancy complications is a novel observation.

During pregnancy, the immunomodulatory effects of steroids are required to maintain a tolerogenic environment to support fetal growth and development. Environmental exposures during pregnancy can elevate concentrations of corticosterone, potentially altering maternal circulating metabolites or crossing the placenta, and affecting fetuses in mice [38, 39]. Serum corticosterone concentrations were significantly elevated in *Pb*-infected compared with mock-infected dams at 8 dpi (**Figure 3G**), suggesting that maternal stress increases during peak parasitemia.

Progesterone signaling is essential to support placental development and fetal growth in mice and humans [40, 41], with concentrations increasing over the course of gestation [40]. Consistent with other infections during pregnancy [24], circulating progesterone concentrations were significantly reduced in *Pb*-infected compared with mock-infected E10 dams (**Figure 3H**). The combination of elevated corticosterone and reduced progesterone likely contributes to immune dysregulation and adverse pregnancy outcomes in *Pb*-infected dams.

### Pb infection during pregnancy does not alter the maternal fecal microbiome

The maternal gut microbiome is crucial in maintaining physiological homeostasis during pregnancy. Whether malaria infection during pregnancy impacts the composition and communities of the maternal gut microbiome is not well defined. We characterized the fecal microbiome of non-pregnant and pregnant mock and *Pb*-infected E10 dams by performing shotgun metagenomic sequencing on free-drop fecal samples at peak parasitemia (8 dpi). The mock and *Pb*-infected dams showed no significant differences among alpha diversity indexes, including richness (number of species) and Shannon diversity at 8 dpi (**Figure 4A**). To assess the relationship between microbial community structure, pregnancy, and *Pb* infection, we calculated Bray-Curtis dissimilarity distances. A Principal Coordinates Analysis (PCoA) of the Bray-Curtis dissimilarity distance showed no separation between *Pb* and mock individuals but revealed distinct microbial communities based on pregnancy status (**Figure 4B**). These findings were further reinforced by a permutational multivariate analysis of variance (PERMANOVA), indicating that pregnancy status (p=0.002) was a stronger driver of differences among samples compared to infection status. Collectively, diversity results highlight that the fecal microbial diversity during *Pb* infection was unaffected during peak parasitemia.

**Figure 4.**
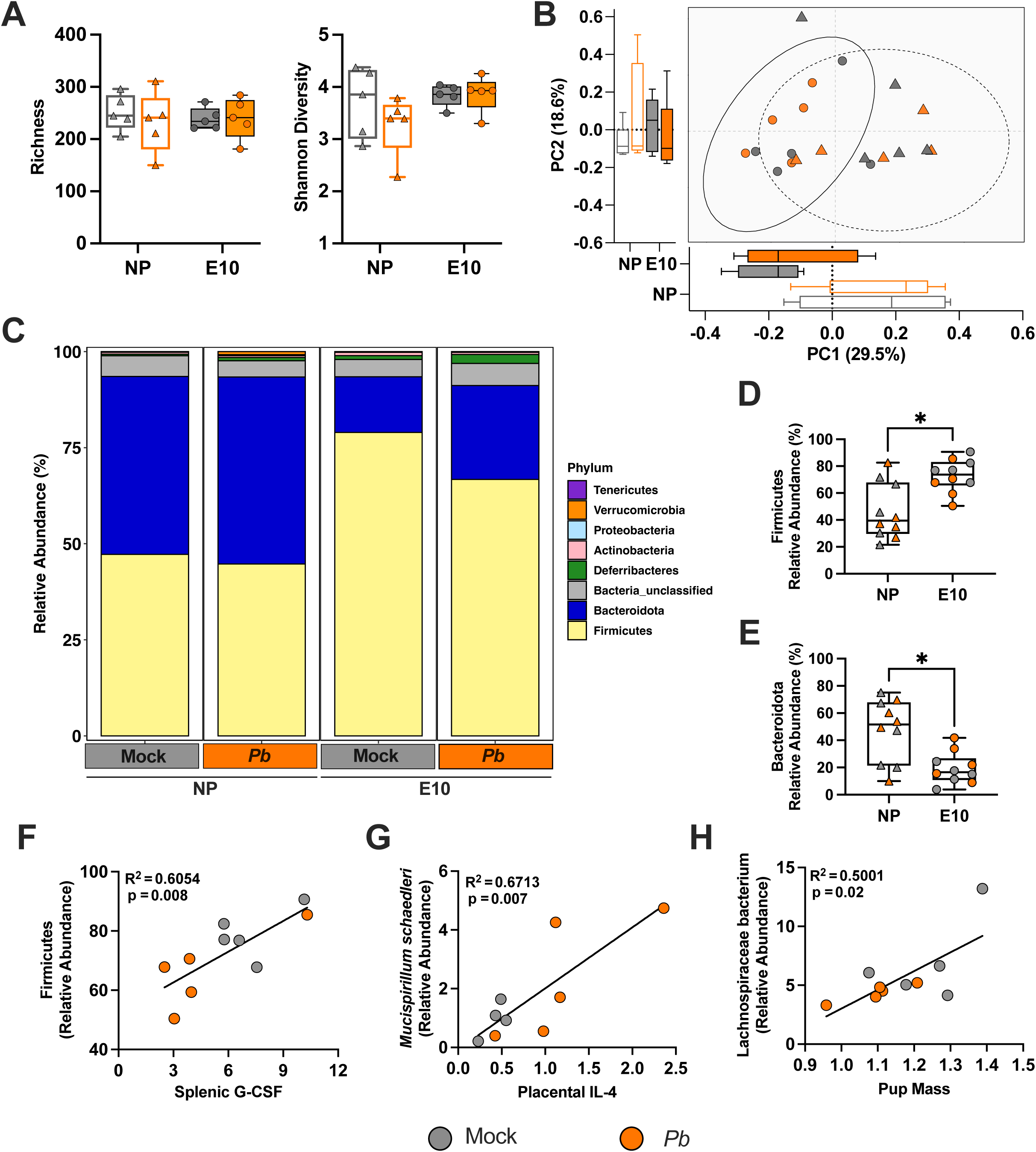
*P. berghei (Pb)* infection during pregnancy causes minimal changes in the maternal fecal microbiome during peak parasitemia. (A) Alpha diversity measures of the observed number of species, or richness (left) and Shannon diversity (right), were calculated and compared between non-pregnant females and embryonic day (E) 10 mock and *Pb*-infected dams at 8 days post-infection (dpi). Pair-wise comparisons were conducted using the Mann-Whitney test. **(B)** PCoA plot of weighted Bray-Curtis dissimilarity distance. Ellipses were constructed using the standard deviation of non-pregnant (NP) females (dashed line) versus pregnant dams (solid line). **(C)** Relative abundances agglomerated at the phylum level, separated by infection and pregnancy status. Comparison of the relative abundance of **(D)** Firmicutes and **(E)** Bacteroidota between NP females and E10 dams. Pair-wise comparisons were conducted using the Mann-Whitney test. Spearman correlation analysis was performed between **(F)** Firmicutes and splenic G-CSF, (G) *Mucispirillum schaedleri* and placental IL-4, and **(H)** Lachnospiraceae and pup mass. A simple linear regression analysis was performed and is displayed. For all graphs, the color designates infection status (Mock: grey; *Pb*: orange), and pregnancy status is categorized by the shape (non-pregnant females: triangles; E10-infected dams: circles). n=5 dams/group across 2 experiments. Asterisks indicate p < 0.05.

Although the diversity metrics suggested minimal differences in *Pb-*infected dams, we proceeded to examine the relative abundance of microbial communities, first at the phylum level. Firmicutes and Bacteroidota were the most abundant phyla observed across both mock and *Pb*-infected non-pregnant and pregnant dams (**Figure 4C**). Differences among both phyla were not observed between mock and *Pb*-infected non-pregnant or pregnant mice, suggesting minimal taxonomic changes of the more abundant taxa after *Pb* infection. Significantly increased Firmicutes (**Figure 4D**) and decreased Bacteroidota (**Figure 4E**), however, were observed in pregnant dams when compared to their non-pregnant counterparts. When examining the taxonomic composition at the species level, unclassified Muribaculaceae and Lachnospiraceae were the most abundant in all mice (**Supplemental Figure 2A**); differences, however, were not observed between mock and *Pb*-infected non-pregnant and pregnant dams for unclassified Muribaculaceae and unclassified Lachnospiraceae. Differential abundance analysis performed by Lefse between *Pb*- and mock-infected pregnant dams identified *Muribaculum*, *Phocaeicola*, *Bacteroides*, and *Akkermansia* to be significantly enriched within the *Pb*-infected while no taxa were significantly enriched within the mock-infected pregnant dams (**Supplemental Figure 2B**). We next examined the association of bacterial taxa with splenic and placental cytokines/chemokines and pregnancy outcomes, including number of viable pups, placental mass, placental efficiency, and pup mass. Numerous taxa correlated with host factors (**Supplemental Table 2**). Firmicutes showed a positive correlation with splenic G-CSF (**Figure 4F**), while *Mucispirillum schaedleri* was positively correlated with placental IL-4 (**Figure 4G**). When examining taxa that may be associated with adverse pregnancy outcomes, Lachnospiraceae bacterium was positively associated with pup mass, with separation between mock and *Pb*-infected dams (**Figure 4H**). Overall, the taxonomic relative abundances showed minimal changes at the phylum and species levels between mock and *Pb*-infected dams; positive correlations, however, were identified between maternal taxa and host factors.

## Discussion

Malaria infection during pregnancy is a major public health concern. Placental malaria, which is a serious complication of malaria in pregnancy, is associated with many adverse outcomes for mothers and infants, ranging from maternal anemia and low birthweight to congenital malaria and miscarriage [165]. A limitation of current placental malaria mouse models is that malaria infection results in severe morbidity and mortality within 7 days post-infection [35, 42]. The heightened lethality could result from the combination of an unnatural route of inoculation and the high dose of iRBCs inoculated [13]. The doses of iRBCs are greater than the approximately 10 sporozoites deposited by a mosquito during bite challenges [43]. Therefore, naïve placental malaria models have been restricted to infection late in pregnancy [42]. More attenuated *Pb* strains have been used in recrudescent and immune models [34, 44, 45], but even these attenuated strains become lethal during pregnancy and must be controlled by drug treatment to remain sublethal. There is no epidemiological evidence that third trimester placental malaria results in a greater burden of disease. Thus, current models may not capture the spectra of malaria infection during pregnancy, as infection risk is not limited to the third trimester in humans.

To address some of these limitations, we developed a placental malaria model using mosquito bite challenge of outbred dams infected during either early or mid-gestation pregnancy. Because the placental malaria burden is greatest in women with limited exposure, naïve models of placental malaria are essential to understanding the mechanisms driving pathogenesis. Compared with iRBC murine models of placental malaria, in which death occurs within 7dpi [35, 46], *Pb* infection by mosquito bite challenge was universally lethal in our model within 9-15 dpi. While E6 dams suffered the greatest morbidity, perinatal mortality occurred in both pregnant cohorts, despite lower numbers of parasites in the uterus and spleen with E6 than E10 infection. Placental malaria was confirmed in our model through two independent methods and while parasites were present in the intervillous space of the placenta, no parasites were transmitted to fetuses during either the first or second trimester. These data highlight that infected RBCs accumulated in the maternal vasculature and never traversed into fetal circulation.

During mid gestation, *Pb* caused significant systemic maternal immune activation that was associated with reduced concentrations of progesterone, elevated concentrations of corticosterone, and intrauterine growth restriction of offspring. This model characterizes the endocrine landscape of placental malaria, highlighting systemic hormonal changes that may contribute to adverse perinatal outcomes. Maternal immune activation is a known driver of pregnancy pathologies, with well characterized connections to adverse fetal outcomes [47, 48]. While there are diverse causes of maternal immune activation, maternal infection, including with pathogens that do not infect the placenta (e.g., influenza H1N1), is a potent driver [24]. In many cases, systemic inflammation, including robust cytokine production in distal sites, can contribute to pregnancy and placental pathologies, including adverse fetal outcomes [24]. While iRBCs were present in the maternal vasculature of the placenta, placental damage and immunopathology, including cytokine secretion, were limited. Systemic production of cytokines, however, was observed, suggesting systemic maternal immune activation may contribute to adverse pregnancy outcomes observed during mid gestation. Neutrophilia and eosinophilia were also observed concurrent with significantly reduced lymphocyte and monocyte fractions. The neutrophil-to-lymphocyte ratio – a marker of distress in mice – also was significantly elevated in E10 infected dams [36]. The neutrophil-to-lymphocyte ratio has not been studied in the context of placental malaria but is associated with malaria infection and parasitemia in non-pregnant mice [49–51]. A higher neutrophil-to-lymphocyte ratio has also been associated with the onset of cerebral malaria in experimental mouse models [52] and hypertensive disorders of pregnancy [53, 54].

The maternal microbiome influences changes in metabolism, hormonal status, and immunological responses related to pregnancy [55]. Microbial dysbiosis of the gut during pregnancy has been associated with complications, including gestational diabetes, preterm birth, diminished intrauterine growth, preeclampsia, and eclampsia, which can be further exacerbated by maternal infection [56, 57]. In inbred non-pregnant mice, the gut microbiota composition is a key factor influencing susceptibility to malaria. Vendor-distinct gut microbe communities, for example, enhance susceptibility or resistance to infection with *P. yoelli* following either i.v. or i.p. injections with iRBCs [58–61]. Analysis of taxonomic composition in previous studies showed that higher abundances of Bacteroidaceae, Prevotellaceae, and Sutterellaceae are associated with increased susceptibility to *P. yoelii* infection in non-pregnant mice, while greater abundances of Clostridiaceae, Erysipelotrichaceae, Lactobacillaceae, and Peptostreptococcaceae are linked to reduced parasite burdens [59, 61]. In addition to *Plasmodium* spp., other parasites such as *Toxoplasma gondii* infection in C57BL/6J mice significantly decreases species diversity, reduces the abundance of Firmicutes and Verrucomicrobia, and increases the abundance of Bacteroidetes [62]. In the context of pregnancy, outbred Swiss Webster pregnant mice infected with *P. chabaudi chabaudi* i.v. using iRBCs exhibited lower parasite burdens and improved fetal and postnatal outcomes with a vendor-specific malaria-resistant fecal microbiota transplant compared to malaria-susceptible feces [63]. Although no differences were observed in alpha or beta diversity in our model, we detected enrichment in the family Bacteroidaceae, particularly within the genera *Bacteroides* and *Phocaeicola*, in *Pb*- infected E10 dams.

There are limitations to this study, including that even with the use of mosquito bite challenge with sporozoites, *P. berghei* infection had uniform lethality in early and mid-gestation, even in outbred mice. While a previously determined average number of mosquitos for infection of CD1 mice was used in our development of this model [43], five mosquitos per dam resulted in several dams that were not productively infected (i.e., did not have detectable parasitemia). We were limited to analyzing placenta pathology, immune responses, and microbial dysbiosis at a single time point during peak parasitemia (8dpi), and future studies should evaluate responses and outcomes at additional times during infection, which can be complicated when working with pregnant animals. Natural infection models have been shown to alter vaccine efficacy in comparison to intravenous inoculation, [43] highlighting the valuable potential of this model for testing antiparasitic drugs, monoclonal antibodies, and vaccines to determine if these treatments mitigate or block placental malaria.

Several factors can influence the susceptibility and severity of clinical manifestations of malaria infection during pregnancy, with key factors including pre-existing immunity, parity, and gestation [2]. Expectant mothers in low or unstable transmission conditions are more susceptible to severe disease due to low acquired immunity, while those in high prevalence regions have acquired protective immunity, resulting in less severe disease [2]. When examining parity, primigravid women were more susceptible to placental malaria than multigravid women in high transmission regions [17, 18] [19]. In assessing the gestational age at the time of infection, susceptibility, prevalence of infection, and parasite density peak in the first half of pregnancy and tend to decrease progressively until delivery [2]. While placental malaria was not gestational age dependent in the current model, effects on pregnancy outcomes were gestational age dependent. Whether parity and pre-existing immunity could alter outcomes of placental malaria in our model requires consideration. In summary, we have developed a tractable model of placental malaria that could be expanded upon and used to safely test therapeutic options, given that pregnant patients are excluded from clinical trials of drugs and vaccines.

## Data availability

All data will be made publicly available upon publication and upon request for peer review. Metagenomic sequencing data set is deposited under BioProject PRJNA1208073.

## Acknowledgements

We thank the faculty and staff at the Johns Hopkins Malaria Research Institute Insectary Core for providing infected and uninfected mosquitoes. We thank the staff of the Johns Hopkins Research Animal Resources for animal care, and the Phenotyping Core for hematology analyses. We are grateful to members of the Davis, Klein, and Thompson labs at Johns Hopkins Bloomberg School of Public Health for their feedback on this work.

## Author contributions

Phebe Ekregbesi, Fidel Zavala, and Sabra Klein conceived the experimental questions contained in the funded pilot award. Phebe Ekregbesi, Brittany Seibert, Maclaine Parish, and Patrick Creisher conducted all mouse work. Yevel Flores- Garcia assisted Phebe Ekregbesi with infection protocols and quantification of parasites. Phebe Ekregbesi, Brittany Seibert, Maclaine Parish, and Jennifer Liu conducted cytokine and steroid assays. Phebe Ekregbesi, Patrick Creisher, Joseph Hoffman, and Yevel Flores-Garacia conducted IVIS analyses. Cory Brayton and Phebe Ekregbesi conducted histology and related analyses. Phebe Ekregbesi, Brittany Seibert, and Maclaine Parish organized all data, conducted all statistical analyses, and created all figures. Brittany Seibert, Maclaine Parish, Fidel Zavala, and Sabra Klein wrote the manuscript, and all authors approved the final draft.

---

Supplemental Figure 1. M**o**squito **bite *Plasmodium berghei* challenge in non- pregnant and pregnant dams. (A)** Kaplan-Meier survival curves for E6 and E10 *Pb*- infected dams. n = 7-12 dams/group across 6 experiments. **(B)** Viability was assessed in fetuses collected at embryonic day 18 by cesarian section in E10 dams. Data are represented as the percentage of viable fetuses out of the total number of fetuses across all litters. Percentages within the solid bar represent viable fetuses while percentages above the striped bar represent non-viable fetuses.

Supplemental Figure 2. F**e**cal **microbial taxonomic classification and differential analysis after malaria infection. (A)** Relative abundances agglomerated at the species level, separated by infection and pregnancy status. **(B)** Differential analysis comparing mock and *Pb*-infected E10 dams. Linear discriminant analysis score was calculated using linear discriminant analysis effect size (LEfSe). The color represents the infection status (Mock: grey; Pb: orange) for which that species was enriched.

**Supplementary Table 1.**
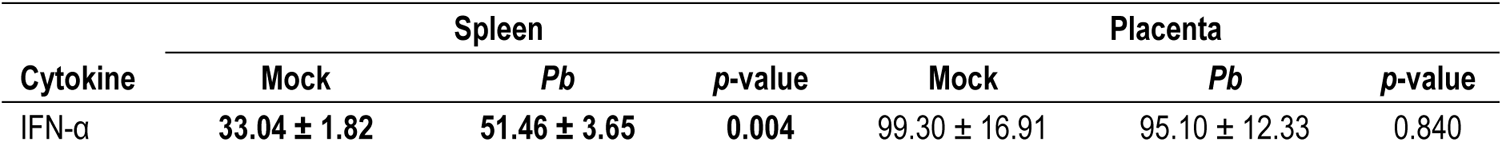
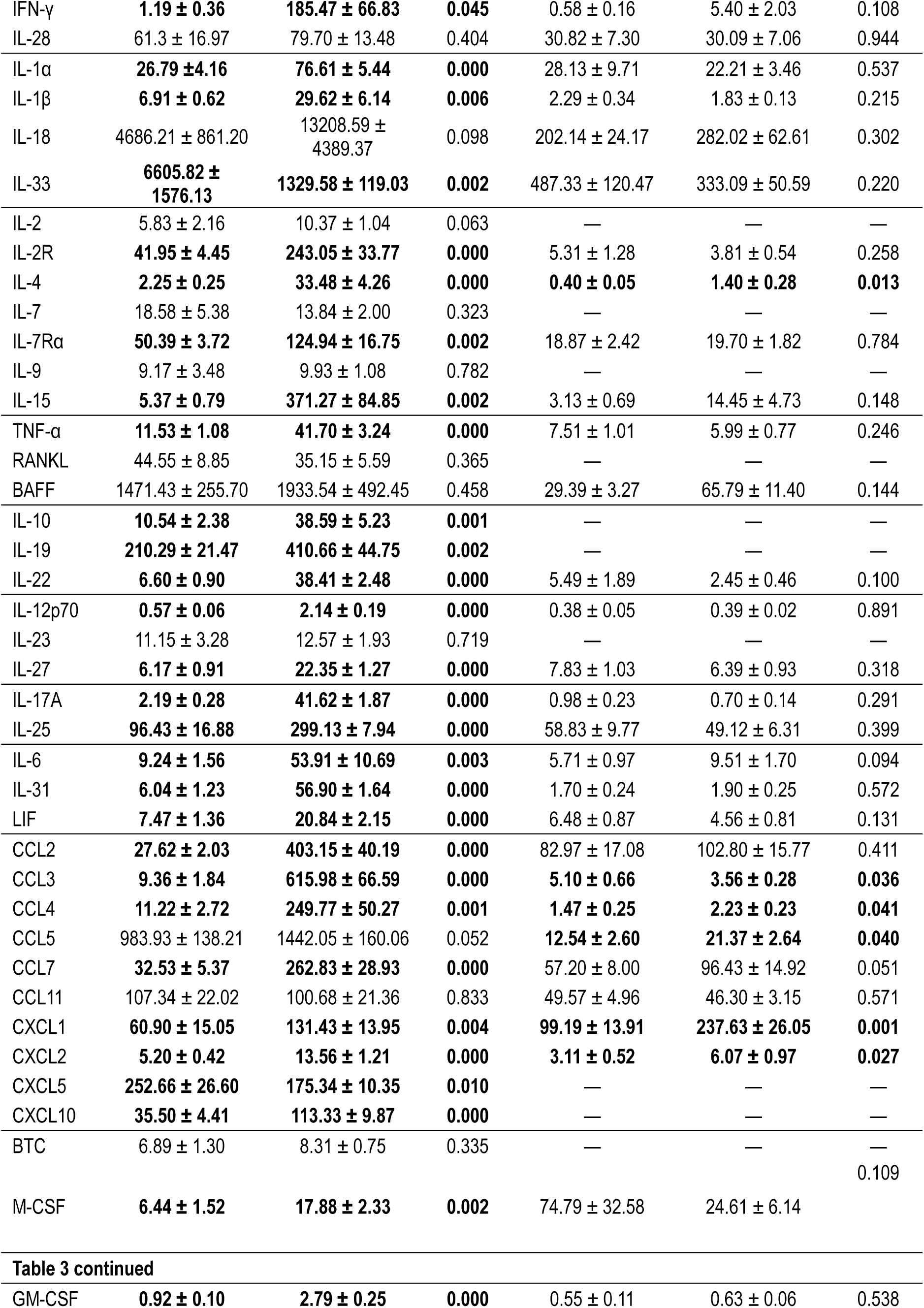
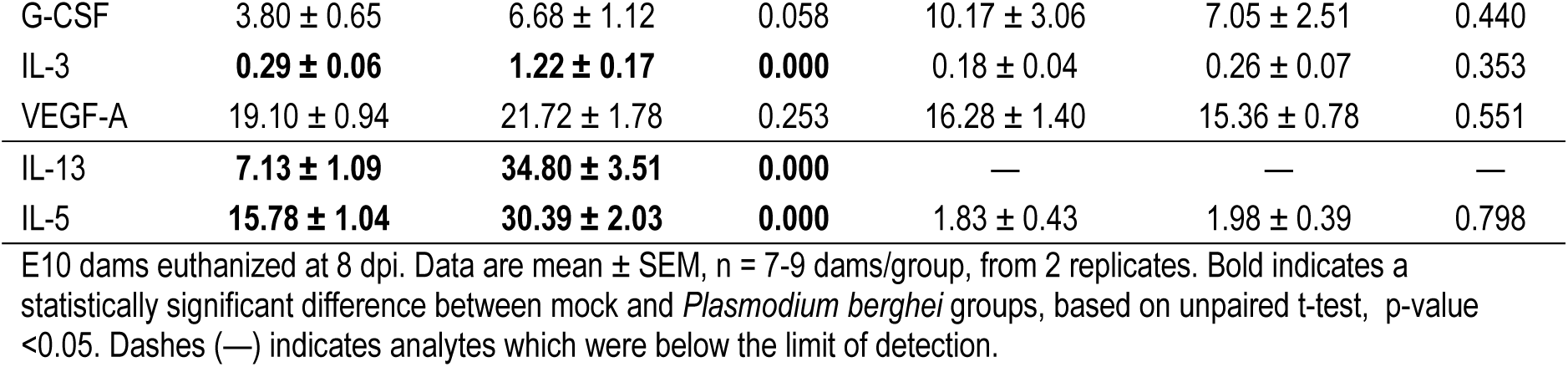
Splenic and placental cytokine concentrations for mock- and *Plasmodium berghei* (*Pb*)-infected E10 dams.

**Supplementary Table 2.** Correlation between bacterial species and pregnancy outcomes in mock- and *Plasmodium berghei* (*Pb*)-infected E10 dams.

| Group | Measure | Taxonomy (Spearman Coefficient, p value) |
| --- | --- | --- |
| Offspring Outcomes | Viable Pups | bacterium 1xD42 87 (0.36, $p=0.037$) |
| | Pup Mass | Lachnospiraceae bacterium (0.54, $p=0.022$) |
| | Placental Mass | Lachnospiraceae unclassified SGB36958 (0.44, $p=0.048$) |
| | Placental Efficiency | Lachnospiraceae bacterium (0.69, $p=0.009$) |
| Placenta Cytokines | IL-4 | *Mucispirillum schaedleri* (0.83, $p=0.007$) |
| Placenta Chemokines | CCL4 | bacterium 1xD42 87 (0.73, $p=0.005$) |
| | CCL5 | *Clostridia* bacterium (0.58, $p=0.046$) |
| | CXCL2 | Oscillospiraceae bacterium (0.54, $p=0.042$)<br>*Acetatifactor muris* (0.70, $p=0.015$) |
| Splenic IFN | IFN alpha | *Clostridia* bacterium (-0.83, $p=0.013$)<br>*Anaerotruncus* sp 1XD42 93 (0.79, $p=0.034$) |
| | IFN gamma | *Candidatus Arthromitus* sp SFB mouse (0.45, $p=0.050$)<br>*Parasutterella excrementihominis* (0.53, $p=0.038$) |
| | IL-28 | *Bacteroides xylanisolvens* (-0.60, $p=0.042$)<br>*Bacteroides thetaiotaomicron* (-0.65, $p=0.046$) |
| Splenic IL-1 | IL-18 | Eubacteriaceae bacterium (-0.10, $p=0.009$) |
| Splenic Common $\gamma$ chain | IL-2 | *Candidatus Arthromitus* sp SFB mouse (0.83, $p=0.006$)<br>*Adlercreutzia caecimuris* (0.77, $p=0.034$)<br>*Parasutterella excrementihominis* (0.50, $p=0.037$) |
| | IL-7R alpha | *Clostridia* bacterium (-0.67, $p=0.041$) |
| | IL-9 | bacterium 1xD42 87 (-0.85, $p=0.032$)<br>*Bacteroides xylanisolvens* (-0.73, $p=0.041$)<br>*Bacteroides thetaiotaomicron* (-0.82, $p=0.041$) |
| Splenic TNF | RANKL | *Parasutterella excrementihominis* (0.61, $p=0.019$) |
| | BAFF | *Parasutterella excrementihominis* (0.79, $p=0.009$) |
| Splenic IL-10 | IL-10 | *Parasutterella excrementihominis* (0.58, $p=0.013$) |
| | IL-19 | bacterium 1xD8 6 (0.62, $p=0.040$) |
| Splenic IL-12 | IL-23 | *Candidatus Arthromitus* sp SFB mouse (0.73, $p=0.046$)<br>*Adlercreutzia caecimuris* (0.75, $p=0.008$) |
| Splenic IL-6 | IL-6 | *Clostridiaceae* bacterium (0.47, $p=0.026$) |
| Splenic Chemokines | CCL4 | *Clostridia* bacterium (-0.68, $p=0.019$) |
| | CCL5 | bacterium 1xD42 87 (0.75, $p=0.021$) |
| | CCL11 | Lachnospiraceae bacterium A2 (0.75, $p=0.007$) |
| | CXCL2 | bacterium 1xD8 6 (0.53, $p=0.031$) |
| | CXCL10 | Clostridia bacterium (-0.65, $p=0.048$) |
| Splenic Growth Factors | Betacellulin | *Parabacteroides distasonis* (-0.77, $p=0.045$)<br>*Bacteroides xylanisolvens* (-0.77, $p=0.010$)<br>*Bacteroides thetaiotaomicron* (-0.82, $p=0.008$) |
| | G-CSF | Muribaculaceae bacterium (-0.65, $p=0.027$)<br>*Duncaniella dubosii* (-0.64, $p=0.041$)<br>Clostridiaceae bacterium (0.62, $p=0.021$)<br>*Adlercreutzia caecimuris* (0.73, $p=0.038$) |
| | M-CSF | Clostridia bacterium (-0.65, $p=0.044$) |
| | VEGF-A | Eubacteriaceae bacterium (0.24, $p=0.027$)<br>Clostridia bacterium (-0.70, $p=0.016$)<br>*Adlercreutzia caecimuris* (0.84, $p=0.019$) |
| Splenic Misc | IL-13 | Clostridia bacterium (-0.73, $p=0.038$) |

